# A multi-omics approach to maize (*Zea mays*) tassel development

**DOI:** 10.64898/2026.01.01.697295

**Authors:** Finn Hartmann, Sandra Mathioni, Atul Kakrana, Blake C. Meyers, Virginia Walbot, Karina van der Linde

**Author notes:** Correspondence: Karina van der Linde.

## Abstract

The development of the male inflorescence in maize (*Zea mays* L.) is a complex and highly regulated process that is essential for reproductive success and yield. To gain a comprehensive understanding of the regulatory programs involved in tassel formation, including the initial specification of anther cell types, the transcriptomes, small RNAs (sRNAs), and proteomes were profiled at four developmental stages (0.5–2.0 cm). The RNAseq analysis indicates dynamic shifts in gene expression underlying the transition from indeterminate meristems to organ initiation and germinal cell determination by 2.0 cm. Complementing these data, sRNA sequencing uncovered 182 microRNAs (miRNAs) with distinct temporal patterns. A core set of 126 miRNAs was expressed throughout tassel development, while others displayed strong stage-specific enrichment. Notably, families of miRNAs that target auxin signaling were dynamically regulated, suggesting fine-tuning of hormone signaling in relation to meristem activity. Later stages were enriched for miR2275 and miR11969, which previously have both been associated with meiocytes, indicating the onset of reproductive sRNA pathways during the early stages of anther differentiation. Together, these datasets provide a broad overview of tassel development and form a backbone to enrich existing RNAseq and single cell RNAseq data sets of specific steps in tassel and initial anther development, while also adding new data on sRNA expression.

## 1 Introduction

Maize, rice, and wheat together account for 30% of the calories consumed by humans. Inflorescence architecture is a key trait for grain production and has been, and still is, a major focus of breeding efforts. During the life cycle of land plants, new organs are generated from distinct meristems containing a set of undifferentiated stem cells. Throughout vegetative growth, organs are primarily generated from the shoot apical meristem (SAM) and the root apical meristem. When the plant switches to reproductive growth, the SAM converts into the inflorescence meristem (IM). The further progression of the IM is species dependent. In *Arabidopsis thaliana*, floral meristems (FMs) form directly on the flanks of the IM. In contrast, grass inflorescences can bear hundreds of spikelets, which are the basic inflorescence units. Each spikelet consists of one or more florets. Rice and wheat have perfect flowers, while maize has separate male and female inflorescences. Maize male flowers are produced on a terminal tassel, while separate female flowers arise on ears in the axils of vegetative leaves. During tassel development, the IM initiates branch meristems (BMs) of indeterminate fate and spikelet pair meristems (SPMs) in parallel (Fig. 1a). The SPMs subsequently divide into two spikelet meristems (SMs). Each SM produces two glumes and initiates the upper FM, then converts itself into the lower FM. Florets contain four concentric whorls that form sequentially from the FM. First, the palea and the lemma are initiated, followed by the lodicule, and three stamen primordia. Finally, carpel primordia are formed. In maize tassel florets, the carpels are aborted early while the stamen primordia continue to differentiate, forming both a terminal anther and a subtending filament. The tip of the primordium undergoes a leaf-like patterning to establish an abaxial side and an adaxial side. These domains are then reorganized into a cross pattern, which allows for outgrowth of the four individual anther lobes between domains. Initially, maize anthers contain cells derived from the meristem layers L1 and L2. The L1-derived (L1-d) epidermis encases the four anther lobes and the centrally located vasculature and connective tissue (L2-d). Within each anther lobe, a subset of L2-d cells specifies into archesporial cells (AR). The remaining, surrounding L2-d cells undergo multiple periclinal divisions and eventually form three distinct layers: the endothecium, the middle layer, and the tapetum. Once all the lobe layers have differentiated, the central AR cells become pollen mother cells (PMCs), which then undergo meiosis I and II followed by a sequence of mitotic divisions ultimately resulting in trinucleate pollen formation.

**Fig 1.**
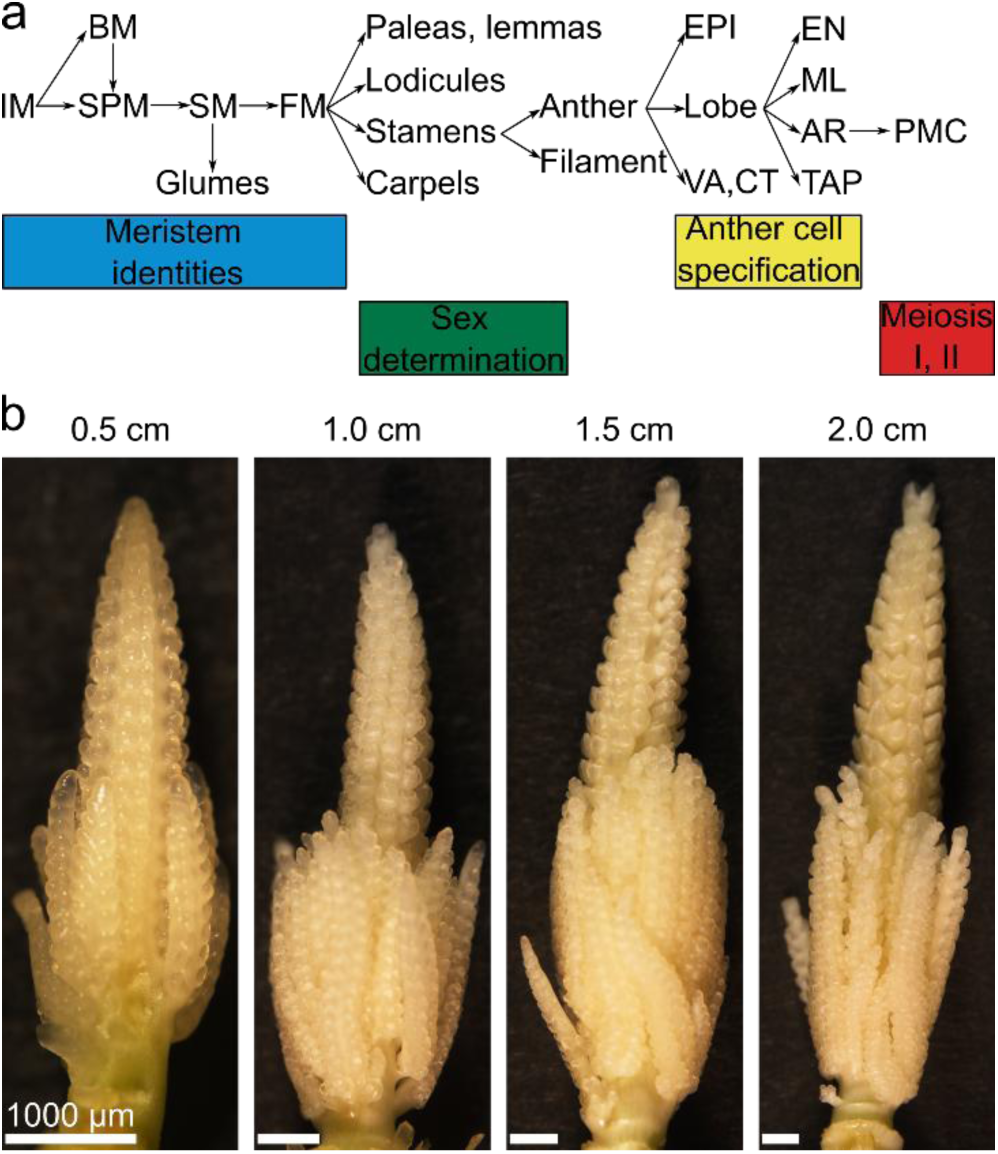
Developmental progression of maize tassels. **a** Male inflorescences development starts with formation of the inflorescence meristem (IM), successively different meristems are formed which finally give rise to the spikelet. In maize the spikelet contains two florets, each containing three anthers. BM, branch meristem; SPM, spikelet pair meristem; SM, spikelet meristem; FM, floral meristem; EPI, epidermis; VA, vasculature; CT, connective tissue; EN, endothecium; ML, middle layer; AR, archesporial cell, TAP, tapetum; PMC, pollen mother cell. **b** Exemplary images of the four different size classes of tassel, which were used in this study.

Decades of research have identified components involved in maize inflorescence development, including the usual suspects, such as phytohormones and transcription factors. A few studies implicate sRNA networks, ligand-receptor interactions, and naturally changing redox conditions. Auxin, among the phytohormones, is required for axillary meristem formation. Mutations in *BIF1* (*BARREN INFLORESCENCE1*) or *BIF4*, which encode AUXIN/INDOLE-3-ACETIC ACID (AUX/IAA) proteins, or in *BIF2* (encoding a regulator of auxin transport), result in a reduced number of branches and spikelets [1–3]. Jasmonic acid (JA) and gibberellins both contribute to sex determination in maize spikelets. In the *ts1* (*tasselseed1*) mutant, JA application rescues stamen development. The lipoxygenase TS1 is most likely involved in the biosynthesis of JA [4]. Gibberellins are responsible for pistil arrest and regulate stamen maturation (Hartwig et al., 2011; Makarevitch et al., 2012; Best et al., 2016). Among the most studied transcription factors in tassel development is RAMOSA1 (RA1), which controls branch meristem determinacy, MADS-box transcription factors, and YABBY transcription factors [5]. The latter are best known for their role in flower development as part of the ABC model [6, 7].

In the early stages of tassel development, microRNA156h (miR156h) and miR172 (encoded by *TS4*) play pivotal roles. miR172 directly controls the expression of *APETALA2* (*AP2*) [8]. Mutant analysis demonstrated that *FUZZY TASSEL*, which encodes a mutated version of a *DCL1* homolog, phenocopies *ts4* [9]. Maize anthers express two different classes of reproductive phased small interfering RNAs (phasiRNAs): premeiotic 21-nt phasiRNAs and 24-nt meiotic phasiRNAs.

Defects in generating these reproductive phasiRNAs have been associated with male sterility in several instances [10–13]. Maize CLE7 (CLV3/EMBRYO-SURROUNDING REGION7), CLE14, and FCP1 (FON2-LIKE CLE PROTEIN1) are small, secreted peptides. FASCIATED EAR2 (FEA2), FEA3, and THICK TASSEL DWARF1 (TD1) are leucine-rich repeat receptor-like kinases (LRR-RLKs). Overproliferation of the IM has been observed in the *td1* and the *fea2* mutants [14, 15]. CLE7 and FCP1 can be sensed by FEA2; however, two different downstream pathways transduce these two peptide signals [16]. In addition, a distinct signaling pathway involves the recognition of FCP1 by FEA3 [17]. The *MAC1* gene encodes a 241 amino acid (aa) protein that is secreted by newly specified archesporial cells as a 218 aa form. Apoplastic MAC1 is required for somatic L2-d cell identity and locally stimulates the periclinal division of neighboring L2-d cells, most probably via the LRR-RLK MSP1 (MULTIPLE SPOROCYTE1) [18]. GRX2, GRX5, and MALE STERILE CONVERTED ANTHER1 (MSCA1) are CC-type glutaredoxins (GRXs) that act redundantly to control the redox state of FEA4, which is a bZIP transcription factor of the TGACG (TGA) motif-binding family [19]. In *fea4* mutants, the IM is enlarged, and the subsequent tassel contains more spikelets and a thicker main rachis [20]. Based on biochemical data, it has been proposed that the oxidized FEA4 dimer binds to DNA and that the reduction of FEA4 by GRXs leads to its monomerization and inactivation. Furthermore, RNAseq analysis of *msca1/grx2/grx5* triple mutants indicated crosstalk with auxin signaling, which is involved in axillary meristem formation during tassel development [1, 19, 21, 22]. The GRX MSCA1 is also essential for AR formation in anthers. Rapid growth and high metabolic demand of this process lead to a naturally occurring oxygen gradient within the four anther lobes. MSCA1 senses the hypoxic signal, resulting in the instruction of the innermost lobe cells to adopt the AR cell fate [23].

In addition to research on individual factors, bulk transcriptomic data has been obtained for the early stages of tassel development [24]. A recent study used single-cell transcriptomics to analyze expression patterns in the IM, SPMs, SMs, and different stages of FMs containing sexual organs [25]. Additionally, proteomic and transcriptomic data of developing anthers have been published [26]. There is a gap, however, bridging the events involved in meristem establishment in the tassel and the initial steps of cell fate determination in anthers. To gain a broad overview of tassel development and provide a backbone for existing datasets that address preceding and following developmental stages, we performed RNAseq on four different tassel size classes. To supplement the existing data and further understand regulatory mechanisms during tassel development, we applied sRNAseq and a proteomic approach. From these data, we chart dynamic changes from the axillary meristem stage to organ formation, including changes in gene expression, translation, and sRNAs.

## 2 Material and Methods

### Plant samples

W23 inbred plants were grown in the Stanford University field or in the Regensburg greenhouse [18, 27]. At approximately 30-33 days post sowing, tassels were dissected and staged by length. For RNA, sRNA, or protein extraction two tassels of the same size class (0.5, 1.0, 1.5, 2.0 cm; +/- 0.1 mm) were pooled per biological replicate and immediately frozen in liquid N2. At the earliest stage (0.5 cm), stamen primordia are present, anthers are readily distinguished by 1.5 cm, and AR specification has occurred by the 2.0 cm tassel stage.

### Tassel imaging

Imaging of tassels at the four above-described stages was performed on a Nikon SZM25 dissecting microscope (Nikon Instruments Inc., Melville, NY, USA).

### RNA extraction and sequencing

Total RNA was extracted using the RNeasy Mini Kit (Qiagen, Germany). Library preparation (300 ng total RNA per sample) and RNAseq were carried out as described in the Illumina “Stranded mRNA Prep Ligation” Reference Guide, the Illumina NextSeq 2000 Sequencing System Guide (Illumina, Inc., San Diego, CA, USA), and the KAPA Library Quantification Kit - Illumina/ABI Prism (Roche Sequencing Solutions, Inc., Pleasanton, CA, USA) at the Genomics Core Facility “KFB - Center of Excellence for Fluorescent Bioanalytics” (University of Regensburg, Regensburg, Bavaria, Germany). Three 100-cycle P2 flow cells per sample were used for sequencing (Illumina, Inc., San Diego, CA, USA). FastQC (v0.11.9) (Babraham Bioinformatics, UK), MultiQC (v1.3) [28] and Trimmomatic software (v0.39) [29] were used to check and remove adapters and reads below 36 nt. Filtered reads were aligned to the maize reference genome B73 RefGen_v4 using HISAT2 [30] with default parameters. Upon quantification of read counts per condition with FeatureCounts [31], expressed genes were filtered (>= 10 counts per gene and condition) and visualized. Upon normalization with variance-stabilizing transformation (vst) of DESeq2 [32], a principal component analysis (PCA) confirmed clustering of biological samples and reproducibility. Variance scores per gene were utilized to analyze expression profiles in a representative subset of 500 most disparate genes between conditions. As the optimal number, *k,* of clusters was achieved by cluster validation methods, K-means clustering (*k* = 4) was performed to partition the data into distinct groups based on similarity, and centroids (cluster cores) were visualized. After a necessary gene-ID conversion with MaizeMine [33], the gene ontology (GO) term enrichment analysis and visualization of biological processes of each cluster was achieved using ShinyGO [34] with default parameters and background genes (filtered for raw read counts > 1). For comparison to earlier developmental tassel stages, we utilized the expressed genes (fragments per kilobase exon per million reads mapped, FPKM ≥ 1) of tassel lengths from 0.3 to 0.7 cm of an existing RNAseq dataset [24]. Similarly, later developmental stages are represented by expressed genes (ON) of premature anther stages (0.15 to 0.4 mm) found in a microarray dataset [26]. All gene IDs were converted to B73 RefGen_v4 identifiers, and the expression values were adjusted accordingly (FPKM).

### Preparation of sRNA and sequencing

Preparation of sRNA and sequencing was performed as described in [35]. A yet unpublished dataset of raw miRNA read counts with a matching experimental design was obtained from the Meyers Lab NGS database [36] under the terms and conditions specified by the repository. All 325 annotated maize miRNAs (18-24 nt) were processed as described for transcriptomics, filtered for raw read counts greater than zero. A tassel-stage specific expression analysis of the 182 remaining miRNAs was performed, as described for transcriptomics, with the exception that miRNAs with a standard deviation of zero of variance between conditions were excluded from further analysis.

### Protein extraction and mass spectrometry analysis

Total protein was extracted as previously described, mixed with SDS-PAGE loading buffer, and proteins were separated (roughly 1 cm into the separation gel) by SDS-PAGE [18]. Lanes were extracted from the gel and submitted to mass spectrometry, as described in Zhang et al (2014) [26]. Sample-specific isobaric labeling was performed as follows: 0.5 cm tassel biol. replicate one, isobaric tag 126; 0.5 cm tassel biol. replicate two, isobaric tag 127N; 1.0 cm tassel biol. replicate one, isobaric tag 127C; 1.0 cm tassel biol. replicate two, isobaric tag 128N; 1.5 cm tassel biol. replicate one, isobaric tag 128C; 1.5 cm tassel biol. replicate two, isobaric tag 129N; 2.0 cm tassel biol. replicate one, isobaric tag 129C; 2.0 cm tassel biol. replicate two, isobaric tag 130N; BLANK, isobaric tag 130C; Pool of all samples to determine ratios, isobaric tag 131. Peptide sequencing was performed as described [26], results were evaluated with the “Byonic” software [37], and peptide fragments were assigned the corresponding protein-isoform IDs of the reference genome B73 RefGen_v3 [38]. With respect to recommended analysis parameters (Bern et al., 2012; Lee et al., 2016; Lu et al., 2020), a peptide-to-protein pipeline was constructed with the following filters: (1) only two missed cleavages were allowed, (2) a False Discovery Rate (FDR) cutoff of less than 1%, and (3) a Byonic Score greater than 200. To infer relative peptide quantities, intensity values from the “Measure” column were used. Peptides were grouped by protein IDs and ranked by “Byonic Score.” The top three ranked peptides were selected for each protein-isoform, mean values were calculated, and replicates merged into a final relative quantification value. Proteins with a value of zero were considered “OFF”, whereas greater than zero was classified as “ON”. Transcript/protein presence was cross-validated using “DeepVenn” [39], after protein-isoform removal and ID conversion using the EnsemblPlants “ID History Converter. In addition, “ON” protein distributions between conditions were illustrated. For further analysis of molecular properties (amino acid sequence and molecular weight, MW), the gene translation file “5b+ gene model set for B73 RefGen v3” was accessed from MaizeGDB. Upon integration, MWs of “ON” proteins per condition were calculated with a publicly available JavaScript [40], and distributions plotted along a capped x-axis (200 kDa) with the R histogram function. Cellular localization motifs, and transmembrane helices (TM) in “ON” proteins were predicted using the algorithms of “TargetP2.0” [41] and “TMHMM2.0” [42]. These predicted features, TM-containing (TM count greater than zero), signal peptide-cont. (SP) and unrecognized proteins were utilized to additionally determine the counts of proteins with only SP motifs, TM domains, or both. Relative feature distributions in total “ON” proteins per condition were calculated and visualized using “GraphPad Prism 8.0.1” (https://www.graphpad.com/). Unique and shared SP-containing proteins per condition were identified using DrawVenn. Lastly, GO term enrichment of biological processes was analyzed and visualized with default parameters and background genes (filtered for raw read counts > 1) using ShinyGO [34].

## 3 Results

To generate overlap with existing bulk sequence data and define distinct stages of male inflorescence development, maize tassels were dissected and examined (Fig. 1b). Tassels measuring 0.5 cm in length still contained an IM region at the tip, while BMs were visible at the tassel base. Both gave rise to SPMs, and SMs had already developed. Glumes were initiated from the SMs in the middle of the main tassel axis. The SMs developed into FMs by the 1.0 cm tassel stage. At a tassel length of 1.5 cm, some side branches still harbor SPMs, while the first spikelets with visible floret structures form on the main tassel branch. These spikelets continued to develop and were fully encapsulated by glumes and lemmas by the 2.0 cm stage. By 2.0 cm tassel length, AR cells were being specified (Kelliher and Walbot, 2012).

### Transcriptomic analysis of tassel development

First, RNA-seq was performed on the four tassel stages. The RNA-seq samples mostly clustered by tassel length in principal component (PC) 1 (Supplementary Fig. 1, Supplementary File 1, 2). A total of 26,408 expressed genes were detected in the four tassel size classes, which indicates a highly complex transcriptome during tassel development. Of the identified transcripts, 23,538 were shared among all tassel sizes (Fig. 2a, Supplementary File 3). Among these transcripts were *AGO18b*, *BIF1-4*, *ZFL2*, *FEA2*, *FEA4, MSCA1*, *GRX2*, *GRX5*, and *TD1*. *MAC1* and *MSP1*, which regulate premeiotic anther development, were also detected in all tassel developmental stages. Even more unexpectedly, *GSL4*, which encodes a GASA (GIBBERELLIC ACID-STIMULATED ARABIDOPSIS) domain protein and was recently described as a resistance factor during *Ustilago maydis* maize infection of seedlings, was among the shared transcripts [43]. Additionally, 56 out of 61 previously identified meiosis-associated genes were expressed at all stages [44]. These results illustrate the importance of “filling in” developmental stages, in this case between tassel architectural development [24, 25] and initial stages of anther development [10, 23, 26, 45]. Transcripts classified as unique to a developmental stage may well be expressed at other as yet untested stages and may play a different role in each case.

**Fig 2.**
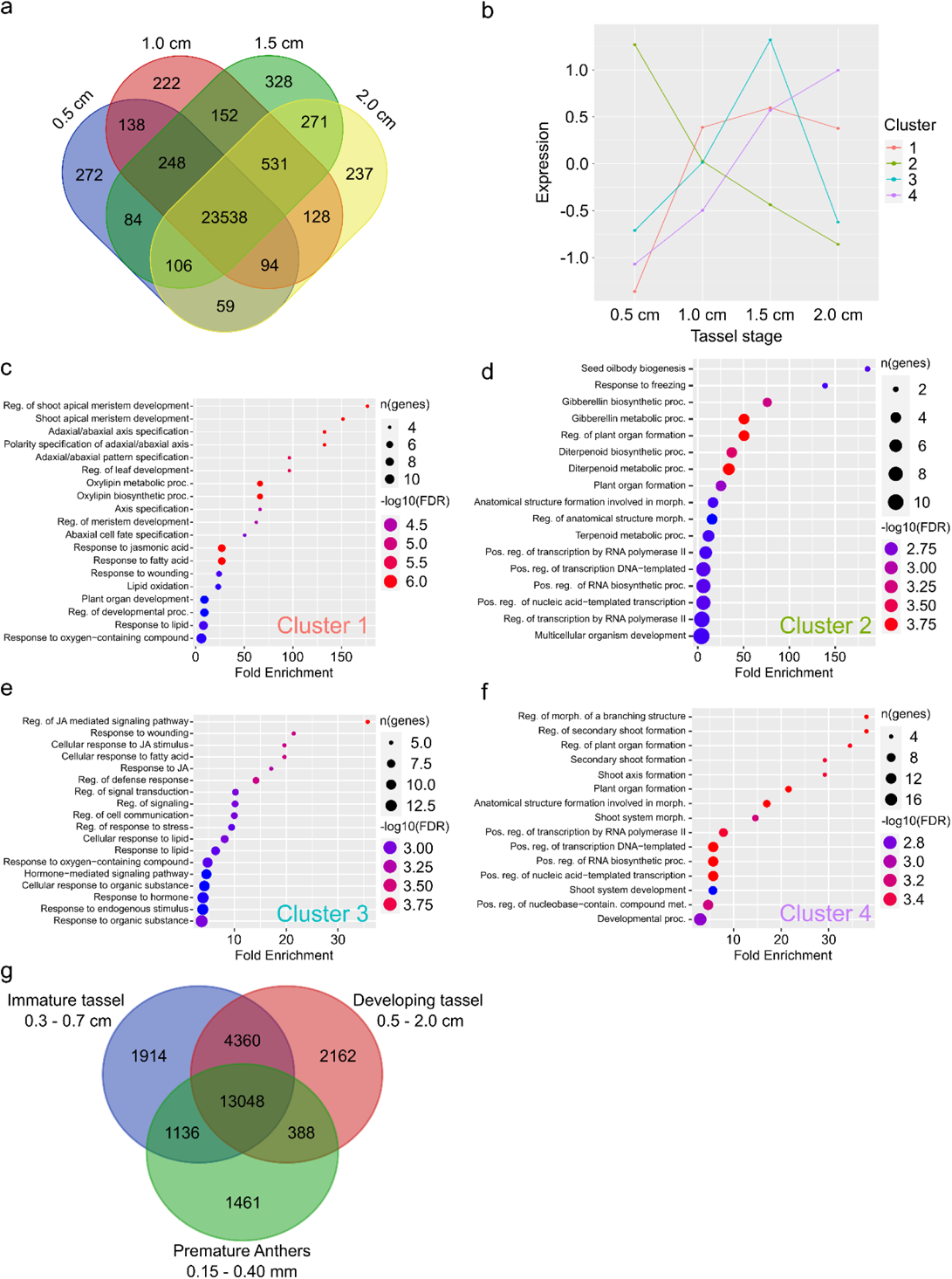
Expression patterns of different tassel developmental stages. **a** Expressed genes (raw, filtered counts of the RNASeq experiment) over four developmental tassel stages (0.5, 1.0, 1.5, and 2.0 cm tassel length), **b** Centroids (mean value of normalized expression values per cluster) plotted over tassel developmental stages, representing the Kmean cluster behavior over time. **c – f** Gene Ontology (GO) analysis of genes for each cluster. GO terms were selected by FalseDiscoveryRate (FDR), sorted by fold enrichment and number of genes are displayed as respective circle radius. **g** Shared and unique transcripts among distinct tassel developmental stages from three datasets: Immature tassel [24], blue; Developing tassel (this study), red; Premature Anthers [26], green.

In 0.5 cm tassels, 272 unique genes were expressed, and 222 genes were exclusively detected in 1 cm tassels (Fig. 2a, Supplementary File 3). 328 genes were expressed only in 1.5 cm tassels. In 2.0 cm tassels, 237 genes were stage-specifically expressed. As expected from prior work, *FCP1* was identified in the 0.5 cm and the 1.0 cm stages, and *RA1* was identified in the stages from 0.5 cm up to 1.5 cm (Fig. 2a, Supplementary File 3). Although the number of shared expressed genes increases during tassel development (138 transcripts shared between 0.5 cm and 1 cm; 152 transcripts shared between 1.0 cm and 1.5 cm; 271 transcripts shared between 1.5 cm and 2.0 cm), these transcripts are 1% or less of the transcriptome shared at all stages. We interpret this result as indicating that at each tassel length sampled, there are cells, tissues, and organs present at slightly different stages, because development is a progression and distinct parts of the tassel are at different points in the trajectory.

To obtain more differentiated data integrating up- and downregulation of genes, K-means clustering was performed on the 500 most disparate genes between conditions (Supplementary Fig. 2, Supplementary File 4). This indicates four distinct expression pattern clusters (Fig. 2b, Supplementary Fig. 3, 4, Supplementary File 4, 5). Clusters 1, 3, and 4 contained genes that exhibited increasing expression from the 0.5 cm stage to the 1.5 cm tassel stage. The expression of genes in clusters 1, 2, and 3 is lower in 2.0 cm tassels than in 1.5 cm tassels. *TS2* was identified in cluster 1, while *TS1* grouped into cluster 3. *GRASSY TILLERS1* (*GT1*) and *FEA3* were associated with cluster 4. Cluster 2 genes showed decreasing expression from the smallest to the largest tassel size (Fig. 2b). These transcripts include *RA3*, *TE1* (*TERMINAL EAR1*), and *UB2* (*UNBRANCHED*2). RA3 and UB2 control tassel branching, and TE1 controls tassel feminization [46–48]. A GO term analysis of biological processes was performed for each cluster. Cluster 1 genes were mainly associated with meristem development or axis specification (Fig. 2c, Supplementary File 6). Cluster 2 GO terms related to gibberellin, diterpenoid processes, and organ formation (Fig. 2d, Supplementary File 6). Conversely, JA-associated GO terms were prevalent in cluster 3 (Fig. 2e, Supplementary File 6). Cluster 4 contained mostly genes linked to organ formation (Fig. 2f, Supplementary File 6).

A previous study analyzed immature tassels ranging from 0.3 to 0.7 cm using RNA-seq [24] . Additionally, an earlier publication investigated the transcriptome of premeiotic anthers from the L2-d stage, before AR development, up to EPI and EN specification [26]. Comparing these two studies with the transcriptomic data obtained here defined 13,048 common transcripts. Of these, 1,914 were uniquely detected in immature tassels, while 4,360 were shared between immature tassels and the tassel stages used in this study including tassels containing anthers. Interestingly, premature anthers shared 1,136 transcripts with immature tassels, but only 388 expressed genes with developing tassels (Fig. 2g, Supplementary File 7). Taken together, the RNA-seq data confirm high transcriptomic activity in developing tassels, which undergoes dynamic changes, as indicated by the differences in the identified GO terms in the four clusters.

### miRNAs across developmental stages

Several miRNAs have been shown to be involved in flower development [for overview see: 49, 50]. To obtain an overall picture of the miRNAs produced during maize tassel development, we subjected the four different tassel stages to sRNA-seq. Overall, this sequencing approach identified 182 miRNAs with detectable expression in at least one of the four conditions. A core set of 126 miRNAs was consistently expressed in all tassel stages, while only a few were stage-specifically detected (Fig. 3a, Supplementary Fig. 5, Supplementary File 8, 9). The highest number of unique miRNAs (18) was detected at the 2 cm developmental stage, and we speculate that these could be associated with initial steps in anther cell fate setting (Supplementary File 11).

**Fig 3.**
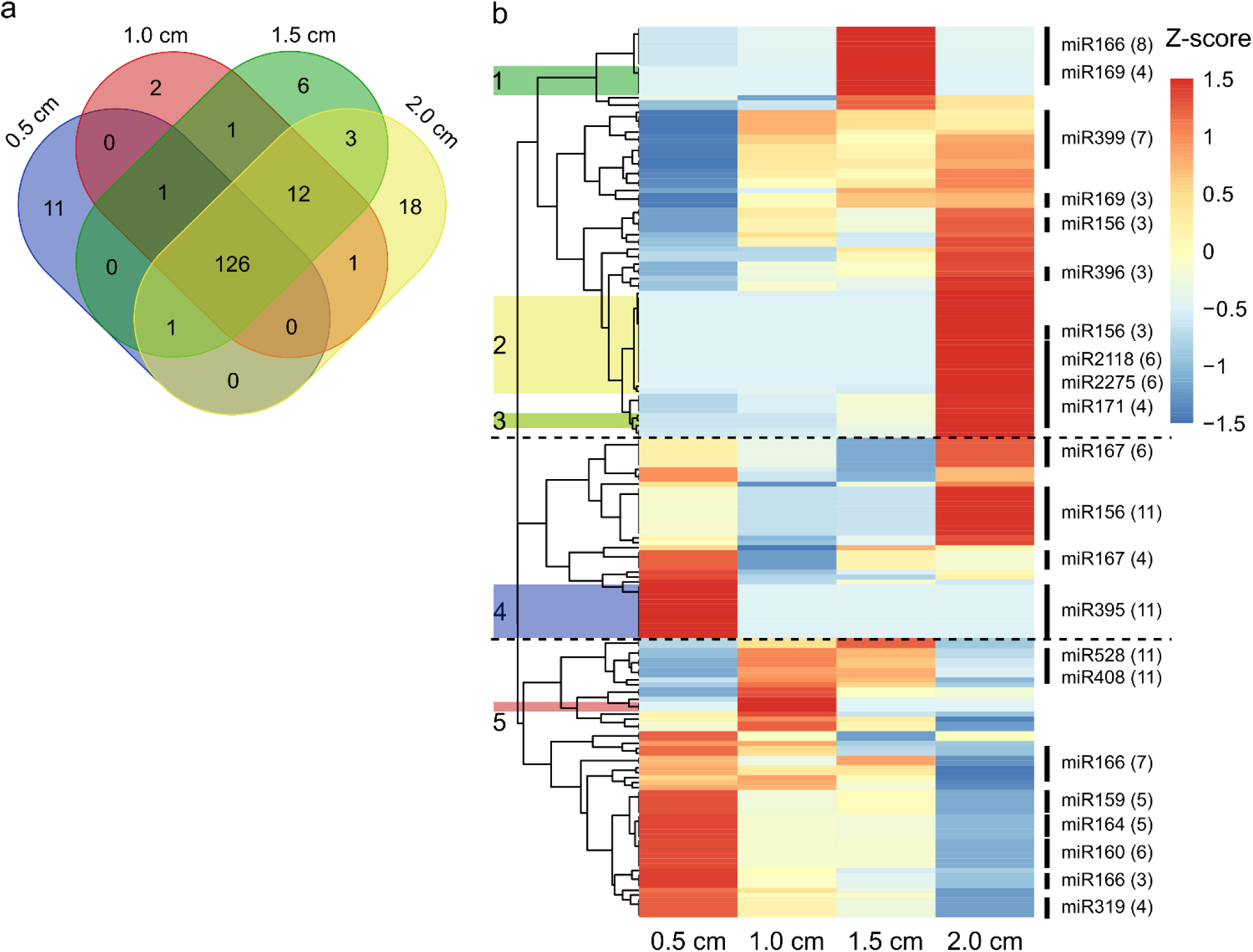
miRNAs in the developing tassels. **a** Abundance of miRNAs (raw, filtered counts > 0) over four developmental tassel stages (0.5, 1.0, 1.5, and 2.0 cm tassel length). **b** miRNA patterns as Z-scores over four tassel stages. Individual clusters identified by hierarchical clustering are separated by dotted lines. miRNA families with three or more members are shown with miRNA-numbers. Colored boxes and numbers indicate miRNAs “Subclusters” found in the Venn-Diagram (Fig 3a).

As with the transcriptomic data, more overlap was identified in the more advanced tassels. To gain a more differentiated view of the data, the normalized reads were standardized, and hierarchical clustering was performed (Fig. 3b, Supplementary File 10, 11). This analysis indicated that distinct abundance patterns of miRNAs can be divided into three main clusters. 1. Enriched in later stages; 2. Lower abundance in the 1.0–1.5 cm stage; 3. Enriched in earlier stages. Within these three distinct clusters, we identified additional subsets of miRNAs. Cluster 1 includes subclusters 1, 2, and 3. miRNAs in subcluster 1 (green) are enriched at the 1.5 cm tassel stage (miRNAs from families 160, 169, and 11696) (Fig. 3b, Supplementary File 10, 11). miR160 and miR169 have both been associated with phytohormones [51, 52]. miRNAs in subcluster 2 (yellow) were mostly detected in 2 cm tassels. They belong to the following miR families: miR2275, 2118, 156, 164, 169, 399, 171, 11969, and 160 (Fig. 3b). Interestingly, this cluster contains two miRNA families (miR2275 and miR11969) that have been identified in meiotic anthers [10, 53]; miR2275 is required for processing of 24-nt phasiRNA precursor transcripts [54]. Recently, pre-meiotic 24-nt phasiRNAs have been identified in maize, and it thus may be relevant that miR2275 is expressed very early in anther development [55]. Other members of the miR2118 family, which is involved in the processing of pre-meiotic 21nt-phasiRNAs, were identified in subcluster 3 (light green), just preceding the temporal period when 21-nt phasiRNA precursor transcripts appear [10]. Subcluster 4 (blue) contains 11 members of the miR395 family and is enriched in the earliest analyzed tassel development stage. Subcluster 5 (red) includes a subset of small. RNAs from the miR167 and miR171 families (Fig. 3b). miR167 has been shown to directly target *Auxin Response Factors* (*ARFs*) in maize [56]. Based on these findings we conclude that, as with the transcriptome, miRNAs are highly dynamically regulated during tassel development and may play a key role in coordinating gene expression.

### Proteomic profiling of tassel development

To complement the transcriptomic and sRNA datasets, we analyzed the proteome of developing maize tassels (0.5–2.0 cm) using mass spectrometry (Fig. 4, Supplementary File 12, 13). After applying stringent filtering criteria (false discovery rate [FDR] <1%, Byonic score >200, and top three peptide fragments per protein ID), we identified a high-confidence set of 4,028 proteins across all developmental stages (Supplementary File 12, 13). This represents a significant increase in the number of identified proteins compared to a previous study on pre-meiotic anthers (Zhang et al., 2014). A comparison with the RNA-seq data indicated that 98.2% of the proteins were associated with transcripts. Only 1.8% of the proteins had no matching transcripts identified (Fig. 4a, Supplementary File 14, 15). We infer that the corresponding transcripts are very short-lived or were expressed at an earlier stage, however, the protein products are sufficiently stable and abundant to be detected. Of the inflorescence development factors previously described and detected by the RNA-seq approach, only FEA3 was identified by the mass spectrometry analysis. It should be noted that GSL4 was also identified by this proteomic approach. Of the identified proteins, 3,928 were present in all four stages of tassel development, while only a few strictly stage-specific proteins were detected (Fig. 4b, Supplementary File 15).

**Fig 4.**
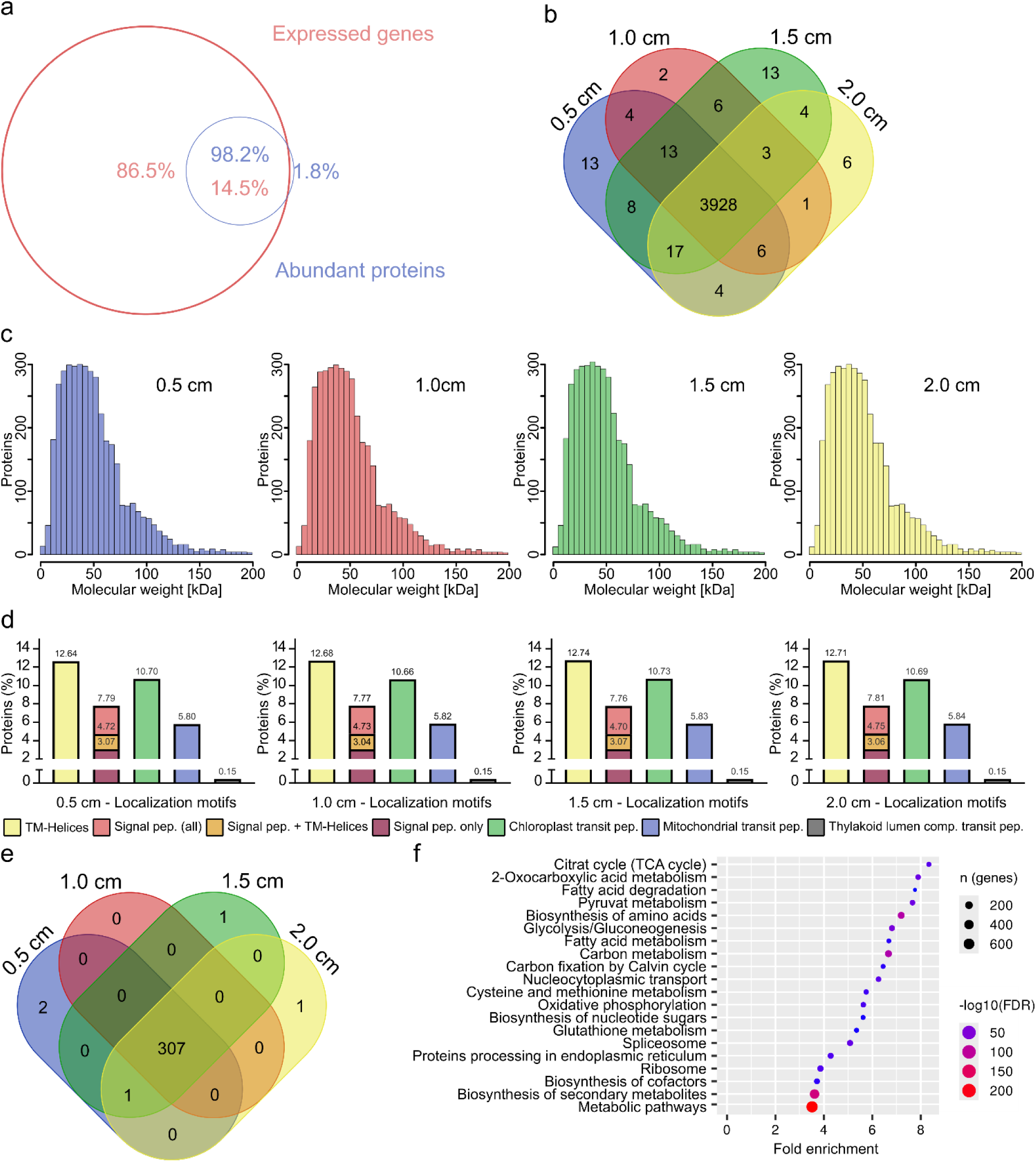
Proteins identified in developing tassels. **a** Comparison of abundant proteins (mean measurement > 0) and expressed genes (raw count >= 10) in developing tassels (0.5 - 2.0 cm tassel length). Percentage of identified proteins for which a transcript was identified by RNAseq are given in red. Percentage of transcripts with a corresponding protein are given in blue. **b** Abundant proteins (blue) shared and unique for four tassel stages. **c** Frequency distributions of molecular weights (MW) in kDa of abundant proteins with a MW limit of 200 kDa. **d** *In silico* predictions of subcellular localization for identified proteins by tassel developmental stage. Predictions are based on presence of N-terminal presequences: Transmembrane (TM)-Helices, secretion signal peptides (pep.), secretion signal peptides with TM-helices, mitochondrial transit peptide (transit pep.), chloroplast transit pep., or thylakoid luminal transit pep. Proteins without target motifs are not displayed. **e** Unique and shared signal peptide-containing proteins in the four different tassel developmental stages. **f** GO term analysis with the top 20 biological processes of all identified proteins. GO terms were sorted by fold enrichment. The number of genes is displayed as respective circle radius, and FDR is indicated by color.

As expected, the molecular weight (MW) distribution of the detected proteins was highly similar across all four stages, with most proteins ranging from 10 to 100 kilodaltons (kDa) (Fig. 4c, Supplementary File 16). Only a few high MW proteins (>150kDa) were detected. To evaluate the potential functional properties of the identified proteins, we predicted their subcellular localization features (Fig. 4d, Supplementary File 17). Across all tassel stages, approximately 12% of proteins contained transmembrane helices and approximately 7-8% carried signal peptides. A fraction of these proteins (approximately 3%) contained both motifs. We identified plastid targeting or mitochondrial transit leader sequences at frequencies of approximately 10% and 6%, respectively. Only a negligible number of proteins were predicted to localize in the thylakoid lumen. The relative proportions of these categories remained stable across developmental stages. We further examined the subset of proteins predicted to enter the secretory pathway (signal peptide-containing proteins). A core set of 307 secretory proteins was shared between all stages, with very few unique proteins per condition (Fig. 4e, Supplementary File 18). The limited stage-specificity of secreted proteins, together with the stable proteome overall, contrasts with the more dynamic transcriptional and sRNA regulatory layers. This could imply that transcript abundance is not a good indicator of protein abundance, and that only highly abundant proteins were detected by the mass spectrometry approach. GO term analysis of all identified proteins indicate that most are involved in plant metabolic pathways and the synthesis of secondary metabolites (Fig. 4f).

## 4 Discussion

In this study, we took a multi-omics approach combining RNA-seq, sRNA-seq, and proteomic analysis to gain a comprehensive understanding of gene expression and protein dynamics during critical stages of maize tassel development. Previous studies have provided transcriptomic atlases of immature tassels and profiled premeiotic anthers [24, 26]. More recently, Sun et al (2024) [25] performed RNA-seq on individual tassel meristems and scRNA-seq on two FM stages. A critical gap in knowledge concerns the transition from tassel architectural definition to the onset of anther cell fate specification. We sampled tassels between 0.5 and 2.0 cm (Fig. 1) to fill in this gap. The overlap of core genes in our analysis with those in previous studies by Zhang et al (2014) [26] and Eveland et al (2014) [24] confirms the robustness of our data and bridges developmental gaps (Fig 2).

The proteomic analysis performed here defined a core set of proteins present across all tassel stages (Fig. 4). The low stage-specificity and high overlap of protein identities contrast sharply with the dynamic transcriptome and miRNAome identified. This discrepancy may imply a lag between transcriptional regulation and protein accumulation or may reflect the limits of mass spectrometry detection. Only 14.5% of all transcripts matched a detected protein, and the enrichment of GO term categories associated with metabolism, as well as a previous comparable proteomic study of anther development, point to the fact that the chosen methodology favors highly abundant proteins [26].

Based on analysis of the RNA-seq (Fig. 2) and sRNA-seq data (Fig. 3), we propose that tassel development occurs in three phases: 1. Transition from vegetative to reproductive growth and formation of various meristems in developing tassels (0.5-1.0 cm stage); 2. Dynamic switch from meristem to organ formation (1.0-1.5 cm stage); 3. Organ identity formation including initiation of cell fate specification in anthers (1.5 cm stage onwards). GO terms associated with transcription, gibberellin, and organ formation declined after the 0.5 cm tassel stage, likely representing the switch from the constantly leaf-forming SAM to IM. Gibberellin is involved in controlling flowering time and the switch from vegetative to reproductive growth [57–60]. Surprisingly, however, GSL4, which has been shown to respond to gibberellin in maize roots, was detected at the transcriptional and proteomic levels at all stages of tassel development [61]. *FCP1* expression was detected in the two earliest stages of tassel development. *FCP1* encodes a CLE-like peptide that is likely sensed by FEA2/3 and regulates SAM homeostasis [16, 17]. TE1 controls leaf initiation, internode elongation, and tassel feminization [48, 62]. Consistent with its previously reported expression in the IM and SPMs, *TE1* was exclusively expressed at the 0.5- and 1.0-cm stages [25].

*UB2*, *UB3*, and *TSH4 (TASSELSHEATH4)* showed the highest expression at the 0.5 cm tassel stage. After that, expression declined, which is consistent with the reported triple mutant phenotype of an unbranched tassel [47]. In turn, miR156 family miRNAs, which represses *SQUAMOSA PROMOTER BINDING PROTEIN-LIKE* (*SPL*) factors, including *UB2, UB3,* and *TSH4*, exhibited high expression at the 2.0 cm stage (Fig. 3) [8, 47, 63]. The *RAMOSA* pathway is central to regulating branching and determinacy of axillary meristems in maize tassels [5, 64–66]. Recently, the direct binding of RA2, which is expressed in all stages, to the promoter of TSH4, which represses *TSH4* expression, was demonstrated [67]. *RA3* and *RA1* were also identified as early-expressed genes (Fig. 2). Taken together, our data support models in which RA3 and SPLs promote branching and meristem boundary formation before being suppressed by an RA-miR156 module during determinacy.

Our data support the model that molecular reprogramming occurs between tassel lengths of 1.0 and 1.5 cm, which coincides with morphological changes in the tassel, such as the termination of BMs, SPMs, and SMs, and the initiation of florets containing stamen primordia. GO terms associated with transcription, shoot apical meristem (SAM) development, and organ initiation declined after the 1.0 cm stage (Fig. 2). Meanwhile, GO categories associated with meristem determinacy, adaxial–abaxial polarity, oxylipin metabolism, and JA biosynthesis peaked at the 1.0 and 1.5 cm stages. The peak in JA biosynthesis-related transcripts at the 1.5 cm stage is preceded by the high expression of oxylipin metabolism-associated genes at the 1.0 cm tassel size. *TS1* and *TS2* expression is highest at the 1.5 cm stage. *TS1* and *TS2* encode a lipoxygenase and a hydroxysteroid dehydrogenase, respectively, which are involved in JA biosynthesis. These two genes regulate the abortion of pistillate structures in the tassel and promote anther maturation [4, 68]. Overall, based on these findings, we propose that *de novo* synthesis of oxylipins, specifically JA, occurs during this developmental stage to enable sex determination in the tassel.

In contrast, the expression of factors involved in several steps of auxin signaling, such as *BIF1*, *BIF2*, *BIF4*, and *SPARSE INFLORESCENCE1* (*SPI1*), was detected at all tassel stages. SPI1, a YUCCA homolog, is involved in auxin production, while BIF2 is a serine/threonine protein kinase that phosphorylates the auxin efflux carrier PIN1a. BIF1 and BIF4 are Aux/IAA proteins that regulate gene expression in an auxin-dependent manner [1, 3, 69]. Auxin signaling is involved in multiple steps of tassel development, including IM patterning, axillary meristem initiation, and floral organ formation, which implies that tight spatiotemporal regulation is required. Analysis of the sRNAseq data show differential enrichment of miR167 and miR160 family members along the different tassel stages (Fig. 3, Supplementary File 10, 11). These miRNA families have been demonstrated to target *ARF*s in plants [56, 70, 71].

The latest stage of tassel development, when the first anthers are differentiating, is marked by the presence of genes linked to development, organ formation, and morphology. Among the strongly expressed genes is *GT1* (*GRASSY TILLERS1*). GT1 is a transcription factor that regulates the abortion of carpels in male florets of maize [72, 73]. Interestingly, miR2275 and miR11969, which are both associated with meiocytes in maize anthers, were enriched at the latest stage (Fig. 3, Supplementary File 10, 11). miR2275 is required for the production of 24-nt meiotic phasiRNAs in anthers [10, 74]. miR11969 was first identified as one of the most highly expressed miRNAs in meiocytes [53]. The recently highlighted presence of 24-nt phasiRNAs at the earliest stages of anthers evaluated, may require miR2275 for processing [55].

## 5 Data availability

This publication contains raw omics data that has been deposited in the NCBI Sequence Read Archive (SRA) (accession number: XXX), and in the Meyerś lab depository.

## 7 Author Contributions

KvdL, SM, and VW performed experiments. Data were analyzed by KvDL, FH, and AK. BM secured research funding and provided supervision. KvdL, FH, and VW wrote the manuscript with input from all authors.

## 9 Supplementary material

**Supplementary figure 1** Principal component analysis of normalized RNAseq counts.

**Supplementary figure 2** Heatmap of expressed genes.

**Supplementary figure 3** Heatmap of K-means clustering.

**Supplementary figure 4** K-means clustering of all expressed genes per cluster.

**Supplementary figure 5** Principal component analysis of miRNAseq counts.

**Supplementary file 1** Raw RNAseq sequencing statistics.

**Supplementary file 2** RNAseq mapping results of HISAT2.

**Supplementary file 3** Overlap and differences in expressed genes.

**Supplementary file 4** Genes of K-means clusters, centroids and correlations.

**Supplementary file 5** Core genes (centroids) per K-means cluster.

**Supplementary file 6** GO terms of biological processes per cluster.

**Supplementary file 7** Comparison of expressed genes.

**Supplementary file 8** miRNA sequencing statistics and raw count table.

**Supplementary file 9** Comparison of miRNAs across stages.

**Supplementary file 10** Normalized miRNAs and filtered Heatmap input.

**Supplementary file 11** Cluster patterns of miRNAs and miR-families.

**Supplementary file 12** Proteomic dataset (exported from Byonic environment).

**Supplementary file 13** Filtered ON and OFF proteins.

**Supplementary file 14** Shared and unique elements between expressed genes and ON proteins.

**Supplementary file 15** Quantification of expressed genes and ON proteins.

**Supplementary file 16** Molecular weight distribution of proteins.

**Supplementary file 17** Predicted localization motifs and transmembrane helices for proteins.

**Supplementary file 18** Shared and unique potentially secreted proteins.

## Acknowledgments

The authors would like to thank Heather Cartwright, the Carnegie Institution Department of Plant Biology Advanced Imaging Facility, and the Kompetenzzentrum Fluoreszente Bioanalytik (KFB) of the University of Regensburg. Additionally, the authors would like to thank Uwe Schwartz (University of Regensburg) and the Computational Systems Biology group of Prof. Dr. Baumbach (University of Hamburg) for their valuable guidance and support in the development of bioinformatic processing pipelines. We also thank Oliver Bear Don’t Walk IV, John Fernandes, Chong Teng, and Leon Kutzner for experimental support and for initial bioinformatics support. Research in the USA was supported by NSF Grant (IOS13- 1339229) to B. C. Meyers and V. Walbot.

