## Supplemental Fig 1-5 for "A multi-omics approach to maize (*Zea mays*) tassel development"

### Supplementary Figures 1-5

#### Article Title: A multi-omics approach to maize (*Zea mays*) tassel development

**Finn Hartmann[1], Sandra Mathioni[2], Atul Kakrana[3], Blake C. Meyers[2,4,5], Virginia Walbot[6], Karina van der Linde[1*]**

[1]Plant Cell Biology, Biochemistry, and Biotechnology, University of Regensburg, Regensburg, Germany

[2]Donald Danforth Plant Science Center, St. Louis, Missouri, USA

[3]Data Science Institute, University of Delaware, Newark, Delaware, USA

[4]Center for Bioinformatics and Computational Biology, University of Delaware, Newark, Delaware, USA

[5]The Genome Center, University of California, Davis, Davis, California, USA

[6]Department of Biology, Stanford University, Stanford, California, USA

***Correspondence:**

Karina van der Linde:

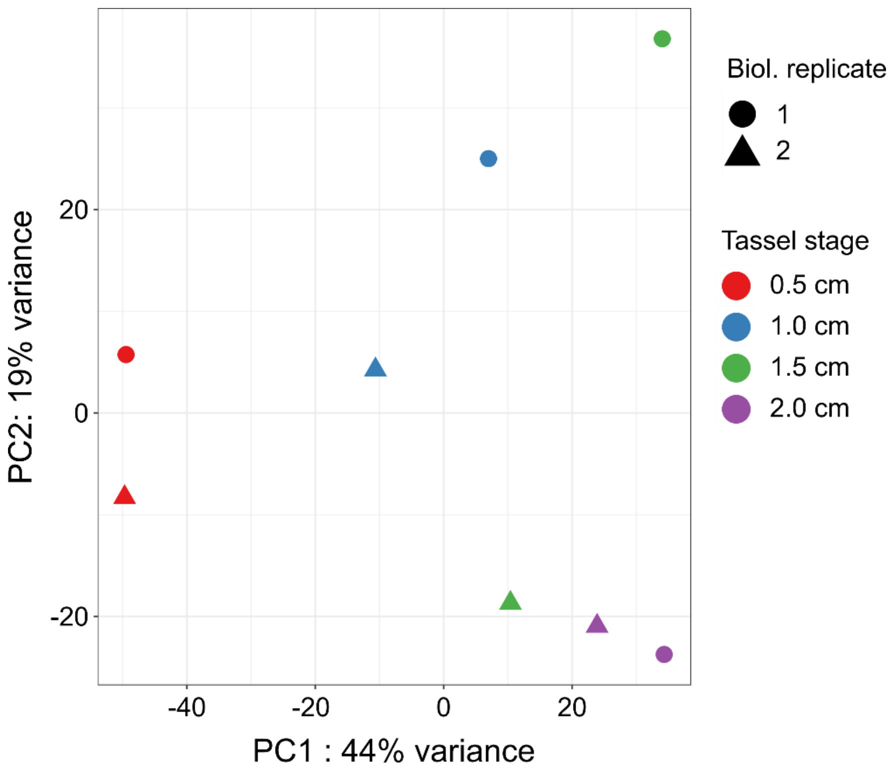


**Supplementary Fig 1 Principal component analysis of normalized RNAseq counts.** Normalized RNAseq counts of two independent biological replicates (biol. replicate 1 = circle, biol. replicate 2 = triangle) were analyzed per tassel stage. Tassel stages are color coded (0.5 cm = red,1.0 cm = blue, 1.5 cm = green, 2.0 cm = purple).

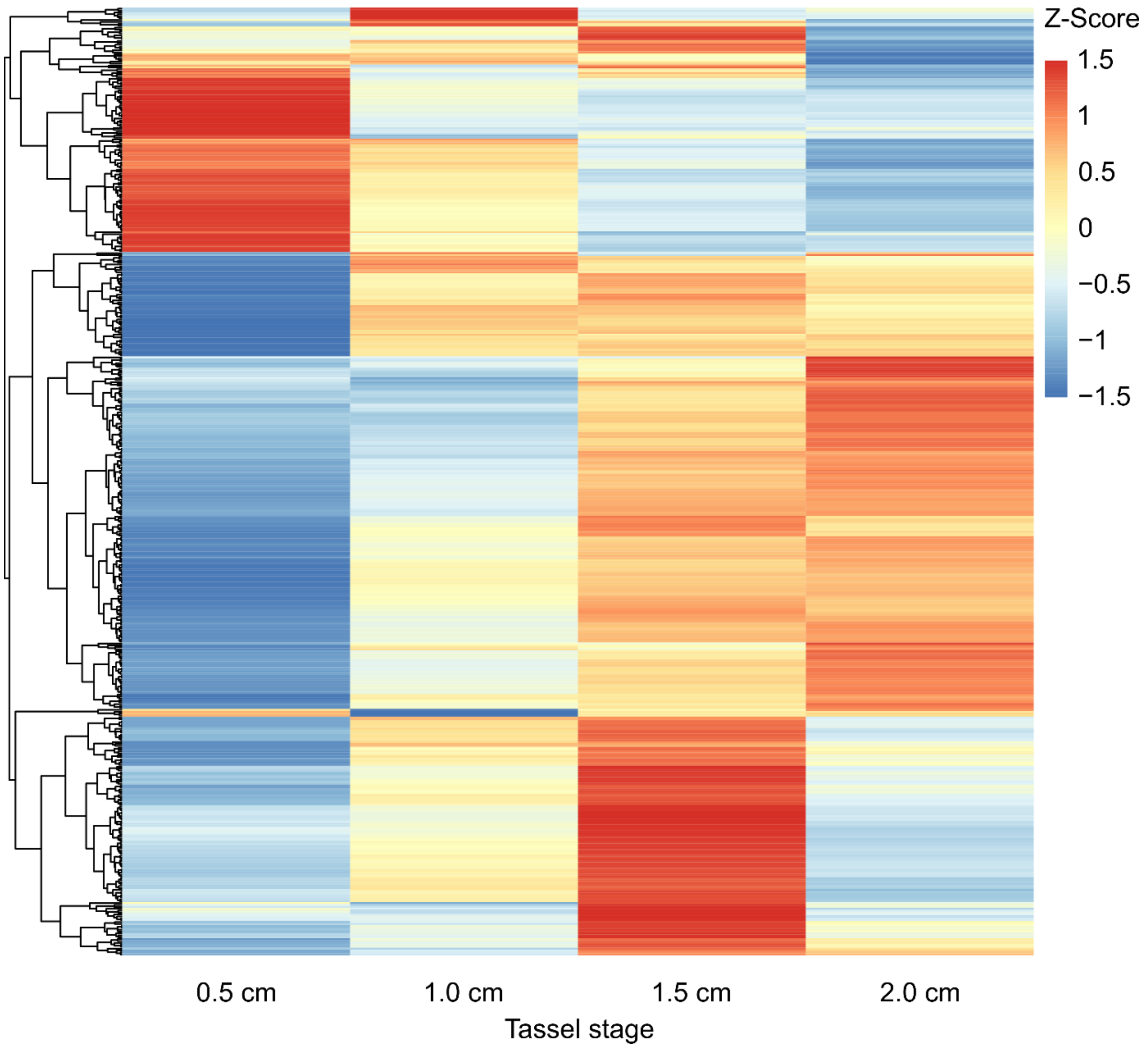


**Supplementary Figure 2 Heatmap of expressed genes.** Z-scores per tassel stage were calculated and hierarchically clustered for each normalized gene expression count.

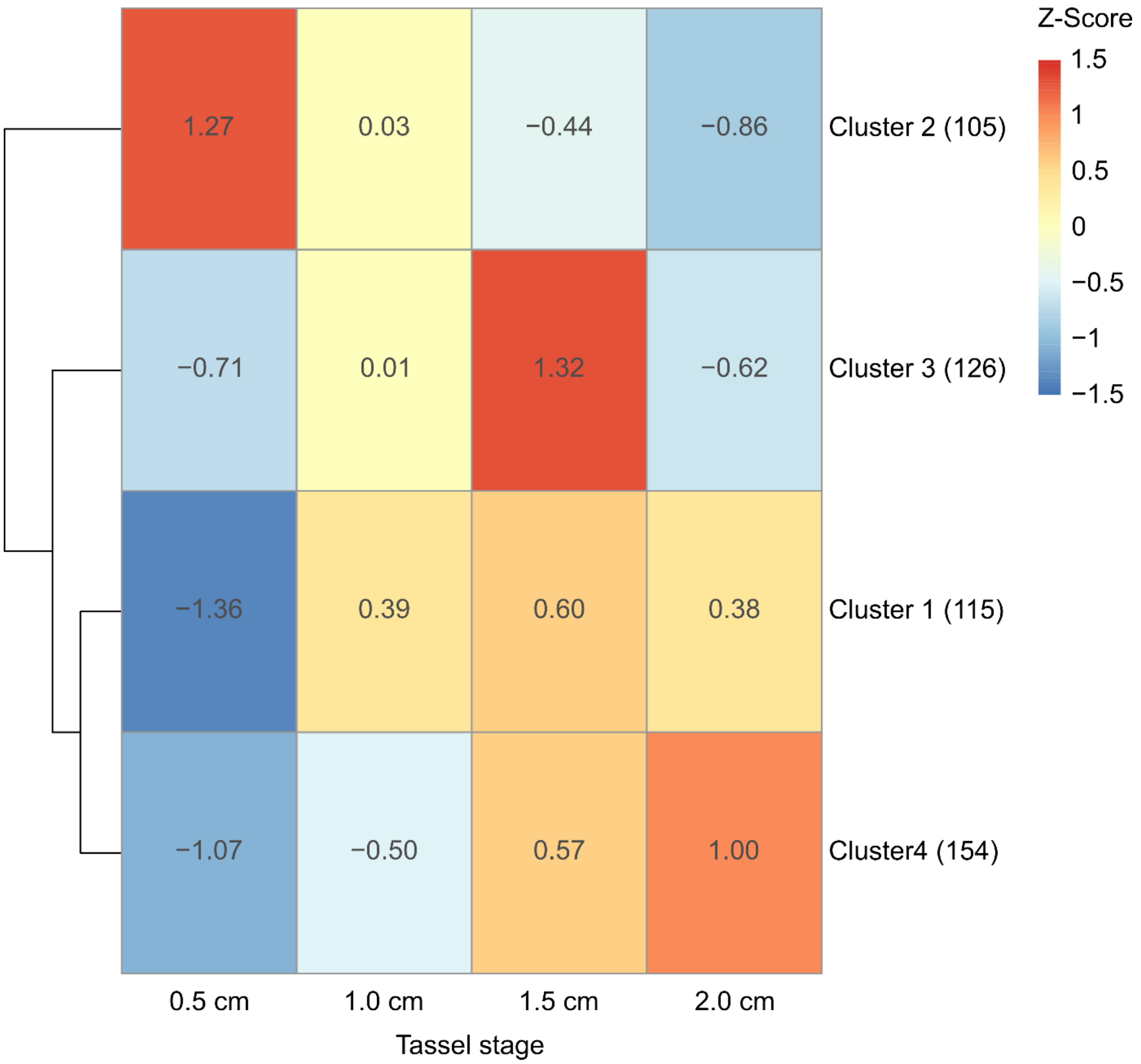


**Supplementary Figure 3 Heatmap of K-means clustering.** The best number of K-means clusters was calculated, and the corresponding Z-scores were clustered hierarchically and by tassel stage

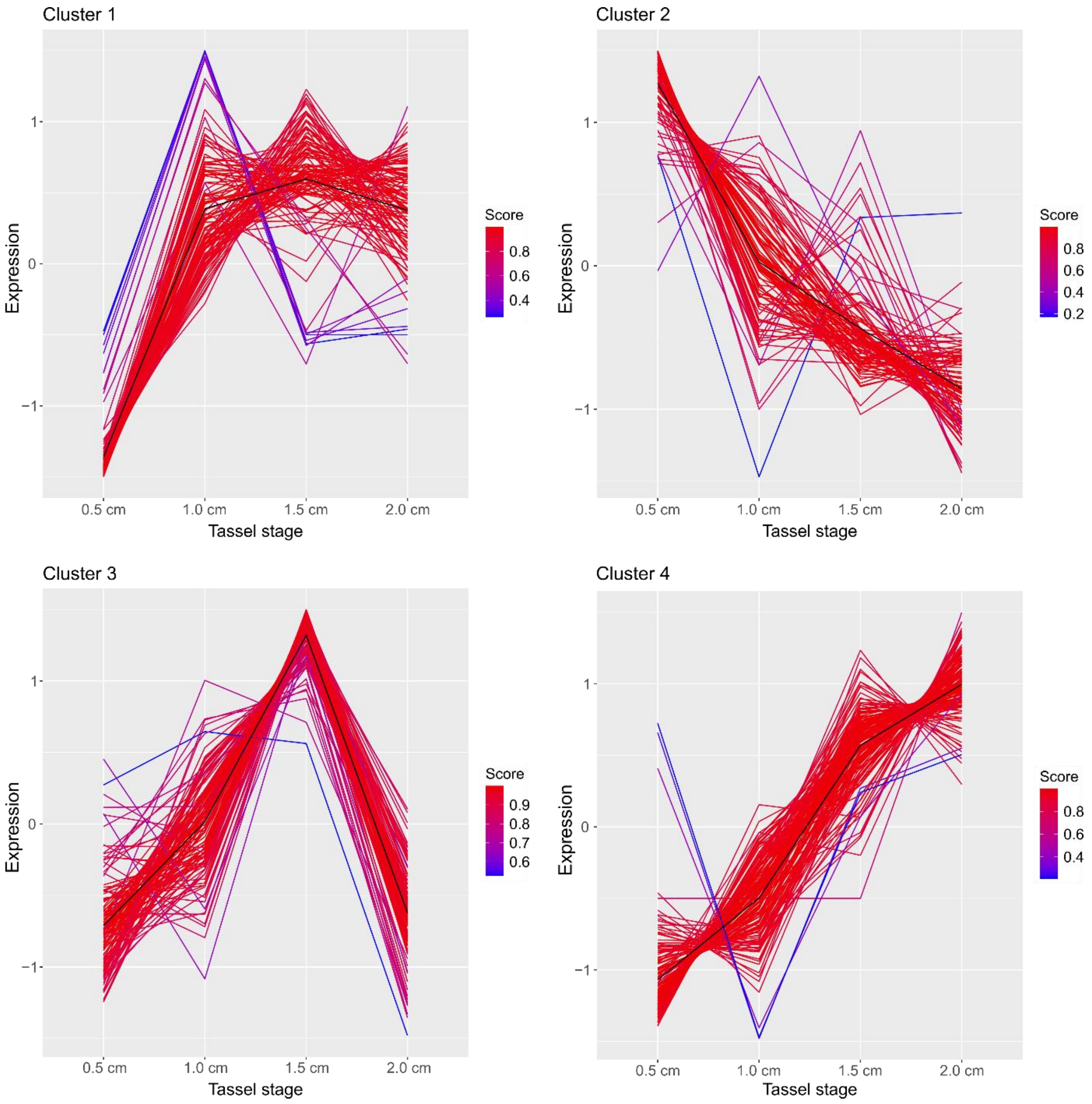


**Supplementary Figure 4 K-means clustering of expressed genes per cluster.** Centroids (mean value of normalized expression values per cluster, black) and all expressed gens per cluster are plotted over tassel stages**.** For each expressed gene the fit score is indicated by color.

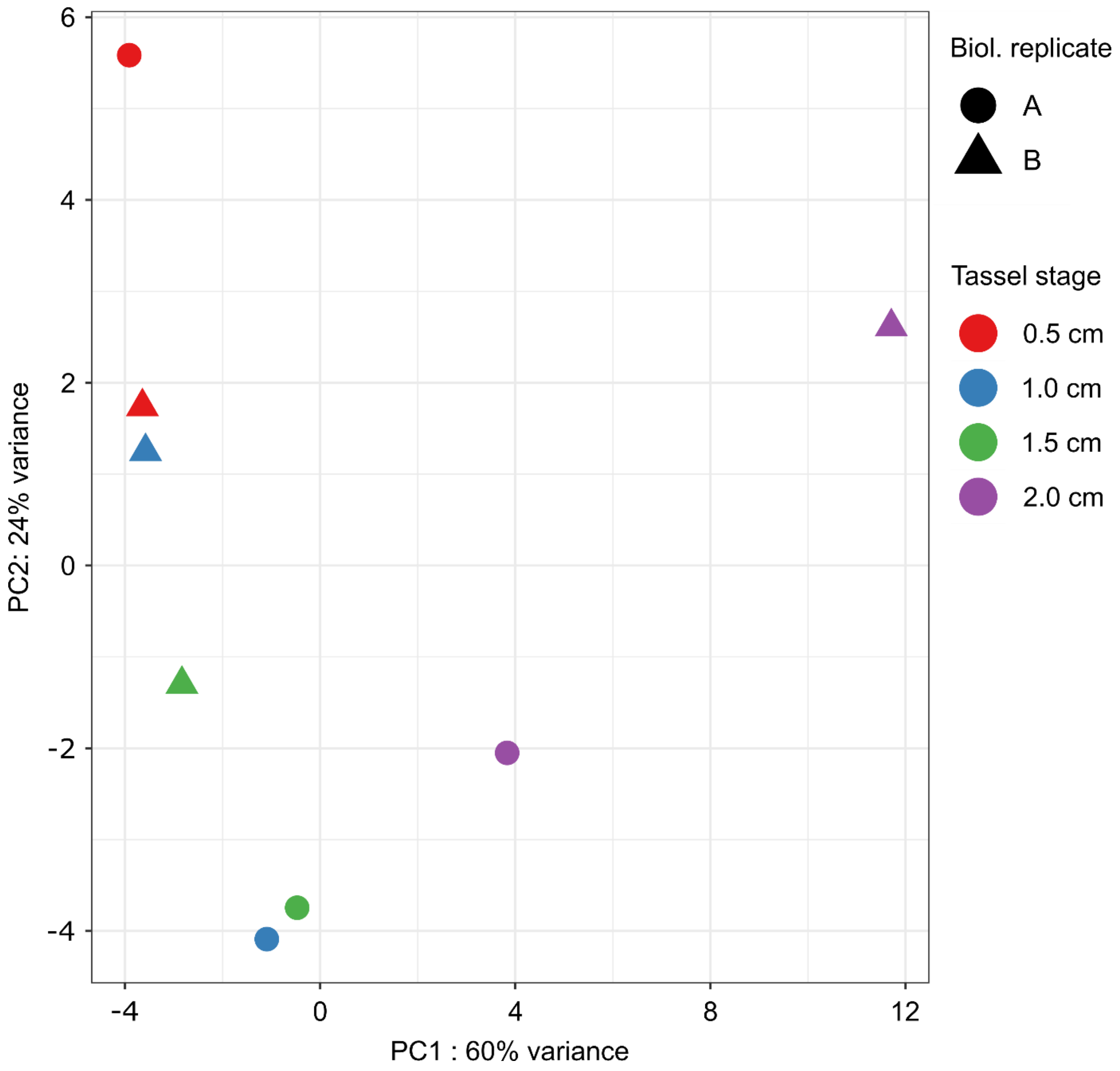


**Supplementary Figure 5 Principal component analysis of miRNAseq counts.** Normalized miRNA counts of two independent biological replicates (biol. replicate 1 = circle, biol. replicate 2 = triangle) were analyzed per tassel stage. Tassel stages are color coded (0.5 cm = red, 1.0 cm = blue, 1.5 cm = green, 2.0 cm = purple).
